# Collective behavior drives diversification across the tree of ray-finned fishes

**DOI:** 10.64898/2026.04.06.716557

**Authors:** Jay Love, Amlan Nayak, Abigail G. Grassick, Ling-Wei Kong, Ella Henry, Michael A. Gil, Ashkaan K. Fahimipour, Matt Pennell, Andrew M. Hein

**Affiliations:** Cornell University; Department of Computational Biology; Ithaca, NY, USA; University of Colorado, Boulder; Department of Ecology & Evolutionary Biology; Boulder, CO, USA; Florida Atlantic University; Department of Biological Sciences; Boca Raton, FL, USA

**Keywords:** Evolution, Collective Behavior

## Abstract

Trait evolution and lineage diversification are thought to be driven by selection on the phenotypes of individuals. However, many important phenotypes are only manifested when individuals interact. Collective behavior is one such phenotype. During collective behavior, interactions among individuals lead to emergent group-level phenotypes that can govern individual fitness. Yet, whether collective behaviors exhibit similar patterns of trait evolution to individual traits and whether these group phenotypes have influenced macroevolutionary diversification is unknown. In ray-finned fishes, the most species-rich of all vertebrate classes, many species exhibit striking displays of collective behavior in the form of collective movement. Here, we use state-of-the-art machine learning methods for text and image analysis to quantify collective movement phenotypes for 2,839 Actinopterygian fishes, comprising nearly 10% of all extant species. We find that collective movement is highly evolutionarily stable in some clades and extremely labile in others, exhibiting over 300 evolutionary transitions. Consistent with theoretical predictions, collective movement is associated with more patchy diets and higher predation pressure. Finally, we find that the evolution of collective movement coincides with an increase in net macroevolutionary diversification rate. Our analyses reveal that collective behavior is deeply intertwined with ecological and evolutionary dynamics across a vast phylogenetic scale, unlocking new questions about the role of collective phenotypes in shaping the diversity of life on Earth.

**Significance:** Collective behavior is thought to contribute to the fitness of individuals that live in groups, but we still do not understand why some species exhibit collective behavior and others do not. We developed machine learning methods to quantify collective phenotypes for thousands of species spanning the phylogeny of ray-finned fishes and found collective behavior to be widespread and consistently correlated with experiencing higher predation pressure and foraging for grouping prey. Lineages that evolve collective movement show a higher net diversification rate. Our results illustrate that collective behavior is foundational to the broad-scale patterns of evolution and diversification in the largest vertebrate class.

## Introduction

Many important behavioral traits cannot be defined outside the context of social interactions (*1*). Such traits challenge standard theory of population and quantitative genetics because an individual’s phenotype is determined not only by genetic and environmental influences, but also by the phenotypes of conspecifics with whom it interacts (*2*, *3*). Some of the most striking patterns of animal behavior observed in nature, collective behaviors (*4*, *5*), fall into this class (*6*). During collective behavior, interactions among individuals within a group can lead to emergent, group-scale phenomena including collective memory (*7*), distributed perception of the environment (*8*), and collective decision-making (*9*, *1*, *10*, *11*), that feed back to determine fitness outcomes of individuals (*12*, *13*, *1*, *14*). How such “collective phenotypes” (*15*) evolve and what kinds of macroevolutionary imprints they may leave on the tree of life has long been debated (*1*, *8*, *13*, *14*, *16–19*). For example, theory (*8*, *16*) suggests that collective phenotypes may evolve more rapidly than individual phenotypes, and recent work has provided some empirical support for this prediction (*18*). Collective behavior also has the potential to impact lineage diversification, for example, by affecting extinction risk (*20*, *21*) or the location and timing of life history events that can induce reproductive isolation among subpopulations (e.g., collective migration [*22*, *23*]). However, due to the challenge of quantifying collective phenotypes across species (*19*), whether these phenomena have shaped broad-scale patterns of trait evolution or diversification within major animal lineages is unknown.

Here, we focus on one of the most widespread forms of collective behavior: collective movement (*6*, *24*). During collective movement, individuals within a group move in ways that are influenced by the positions and actions of the group-mates around them (*25*, *26*), leading to correlated motion at the group scale (*7*). Familiar forms of collective movement occur as murmurations of starlings (*26*), coordinated traffic flow in ant foraging columns (*27*), and synchronized motion of fish schools (*28*, *29*), but interspecific variation in collective movement across the diversity of life has not been well-characterized (*19*). Moreover, the broad-scale macroevolutionary patterns of collective movement have not been well-studied in any animal lineage. Current evidence leads to a number of contrasting predictions about how collective movement may evolve. On the one hand, perceiving and processing stimuli produced by social partners is thought to require specialized sensory adaptations (*30*, *31*) and involves dedicated brain circuitry (*32*, *33*), suggesting that coordinated changes in physiology may be required to enable the evolution of collective movement. Under this view, one might expect collective movement to have evolved rarely. On the other hand, artificial selection and hybridization experiments suggest that, within a species, the propensity to move collectively as well as the quantitative features of collective movement can change significantly within even a few generations (*34*, *35*). This implies that different ecological conditions selecting for or against collective movement could potentially lead to frequent gain or loss of this trait. Consistent with this, the tendency to move collectively with conspecifics varies across subpopulations in zebrafish (*36*), Mexican tetra (*30*), guppies (*37*), and sticklebacks (*31*), and among closely related species of butterflyfishes (*38*). These contrasting lines of evidence and the lack of large-scale comparative analyses of collective movement leave many open questions about how this collective phenotype evolves and how it may have impacted historic evolution and diversification of animals.

We investigate the causes and consequences of variation in collective movement in the Actinopterygians – the ray-finned fishes. Members of this clade, a lineage containing roughly half of all vertebrate species, are known to vary in the degree to which they exhibit collective movement. By quantifying collective phenotypes for species spanning the Actinopterygian phylogeny, we characterized macroevolutionary dynamics of this trait, allowing us to ask whether the evolution of collective movement has been broadly shaped by shared ecological factors and whether its evolution has, in turn, shaped the diversification of the vertebrates.

## Results

### Collective movement is evolutionarily stable in some fish lineages and labile in others

We first sought to characterize broad-scale patterns of variation in collective movement in Actinopterygian fishes. In the fish natural history literature, collective movement is often described as a binary trait (e.g., species are described as “schooling present”(*39*), or “solitary species”). We thus began by treating collective movement as a binary trait in this fashion (below, we extend our analysis to continuous measure of collective movement). Each species was classified as phenotypically *collective* if members of that species are known to move in groups or as *solitary* if they are not (see materials and methods). To inform these phenotypic classifications, we used FishBase (v. 04/2025, [*40*]), a public database containing detailed information on the behavior of fish species and records for more than 33,000 taxa.

In the FishBase database, any information pertaining to collective behavior is contained within blocks of text describing a species’ behavior and ecology; importantly, the level of detail, terminology, and information included in these text blocks varies substantially by species. Following a series of filtration steps to identify species records which contain information related to collective movement (see materials and methods), we applied a few-shot text classifier consisting of a text transformer with a classifier head (*41*, *42*) (Fig. 1a; See materials and methods). This approach uses text descriptions as “input data” to infer each species’ true phenotype, and provides a corresponding measure of uncertainty about this phenotype. This pipeline allowed us to produce standardized, repeatable classification outputs and to account for variable language and ambiguity within the text records. To train and test this model, we manually verified behavioral phenotypes for a subset of the data (11.5% of overall sample, 328 species; See materials and methods) using extensive independent searches of primary and grey literature. After training the model with 50% of these ground-truth species records, the classifier achieved 98% accuracy, 98% precision, 98% recall, and 98% F1 on the test set (i.e., samples not used during training, See materials and methods; Supplementary Information [SI] Section S7). We applied this trained model to the remaining species records, yielding a final dataset containing adult phenotypes (“solitary” or “collective”) and associated confidence scores for 2,839 species spanning 272 families (See materials and methods).

**Figure 1.**
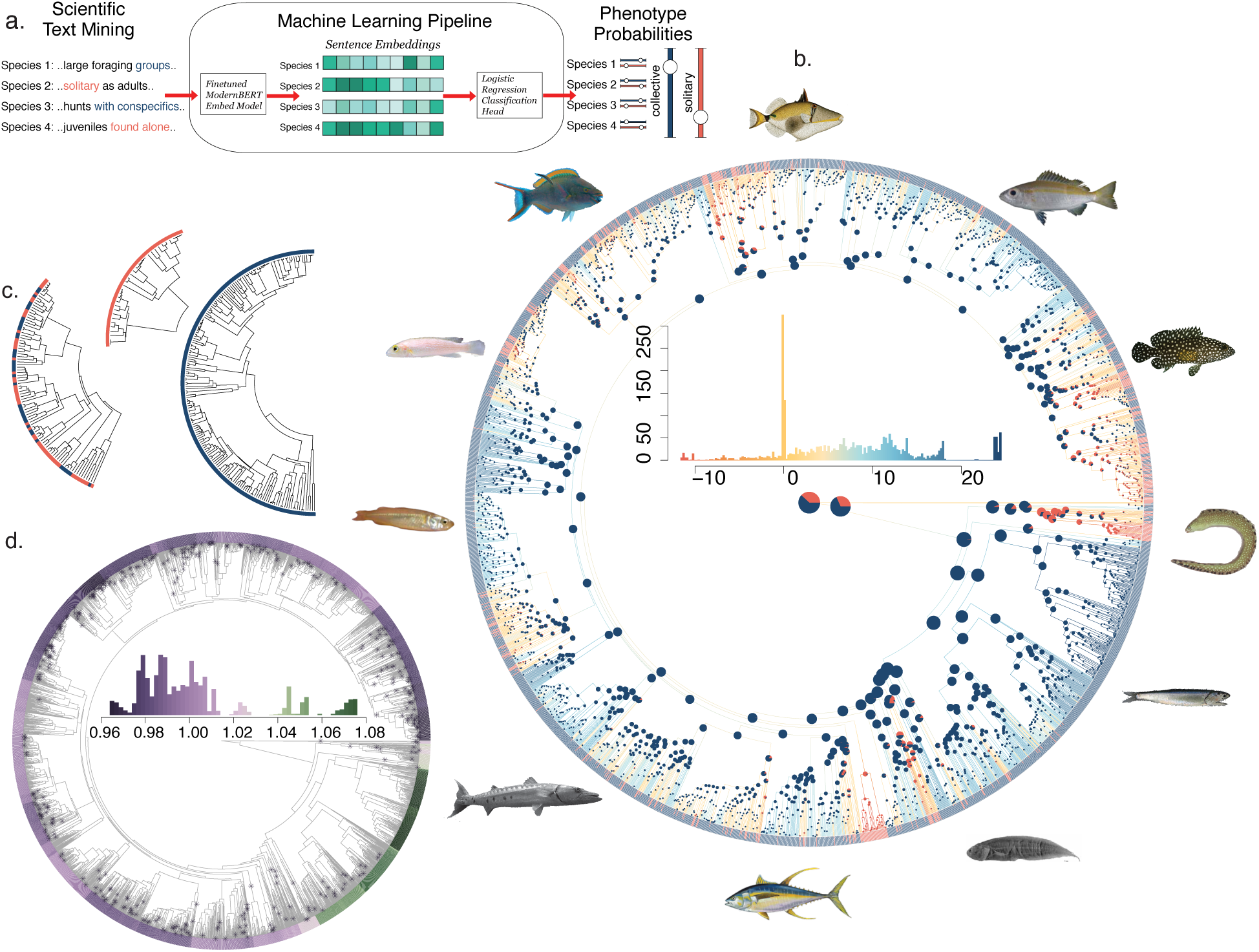
Collective movement is broadly distributed across the Actinopterygian phylogeny. (a) Text classification pipeline to generate phenotype assignments and probabilities. (b) Discrete collective phenotypes (filled circles in tree, blue is *collective* and orange is *solitary*) and distribution of continuous latent trait values, *c* (line colors in tree. Scale shown in inset histogram) show a heterogeneous phylogenetic distribution. (c) Some lineages are composed of a mixture of solitary (red) and collective (blue) species (left subtree). Others are composed of purely solitary (middle subtree) or purely social (right subtree) species. (d) The 388 state transition events, inferred as crossings of the threshold at *c* = 0, are also unevenly distributed (asterisks), with some clades showing few transitions (high stability clades in green), and others showing many transitions (low stability clades in dark purple). Trait evolutionary *stability* is quantified at the species level the quotient of the average phylogenetic distance to all inferred (by the threshold model) threshold crossings and the average phylogenetic distance to all nodes (colors as in inset histogram; SI Section 1).

Within our final dataset, 79% of species exhibit collective movement as adults. Mapping the discrete trait values to the tips of a published phylogenetic tree of Actinopterygian fishes (*43*) pruned to our sample (See materials and methods) shows that collective movement is broadly distributed across the phylogeny (Fig 1b). Comparing clades of similar branch length, many show low within-clade trait diversity, while others show high within-clade diversity (Fig. 1c; SI Section 1, Fig. S1a). This pattern stands in contrast to expectations for a binary trait evolving under a standard Markov process with constant transition rates, but is consistent with evolution of a trait via a threshold process (*44*) (i.e., the trait is underpinned by a more continuous genetic basis but is observed as a discrete trait (*45*, *46*); SI Section 2). Evolution of such a trait can be described by a model originally developed by Sewall Wright, known as the threshold model, which assumes that a continuous latent variable underlies discrete trait values (*45*, *47*). We refer to this latent variable as the “collectivity”, *c*. Positive values of *c* correspond to the collective phenotype and negative values correspond to the solitary phenotype.

By fitting a threshold model to the binary trait data (using the discrete approximation [*48*]) we estimated *c* at all extant tip and internal nodes in the tree (latent extant state and ancestral state reconstruction, See materials and methods). This analysis revealed 388 independent origins and reversions of collective movement, defined as crossings of the threshold value of ! = 0 (Fig. 1b,c). Notably, these threshold crossings are distributed non-uniformly across the phylogeny. The distribution of tip values of *c* indicates many species with *c* far from threshold (Fig. 1b inset). These species are found in clades with relatively low within-clade discrete trait diversity described above (Fig 1b inset, right and left tails; Fig. S1b). At the same time, many species have values of *c* near threshold and are found in distinct clades with high within-clade discrete trait diversity and high incidence of threshold crossings (Fig. 1b inset, central peak; Fig. 1d inset; SI 1, Fig S1b,c). As one might expect, *c* is strongly correlated with the quotient of the average phylogenetic distance to all inferred (by the threshold model) threshold crossings and the average phylogenetic distance to all nodes, a value which we refer to as *stability* (linear regression, DF = 2838, R^2^ = 0.56, *p* = 2.2 × 10^−16^), SI Section S1); species with |*c*| >> 0 tend to be phylogenetically distant from species with the opposite phenotype (SI Section S1).

A corollary of the threshold model is that species with mean values of *c* near the threshold should exhibit mixed phenotypes because some individuals in the population have values of *c* above threshold and others will have values below threshold (*47*). To test this, we focused on a subset of 71 species for which extensive survey data from field studies allowed us to compute a measure of grouping propensity, *g*, for each species, defined as the fraction of field observations in which individuals of a given species were found in groups (See materials and methods). Consistent with predictions for a threshold trait, *g* is strongly positively correlated with *c*, and species with values of *c* near the threshold, *c =* 0, exhibit a mixture of collective movement and solitary movement behavior (logistic regression, DF = 69, Z = 3.24, *p* = 0.0012, Fig. 2a).

**Figure 2.**
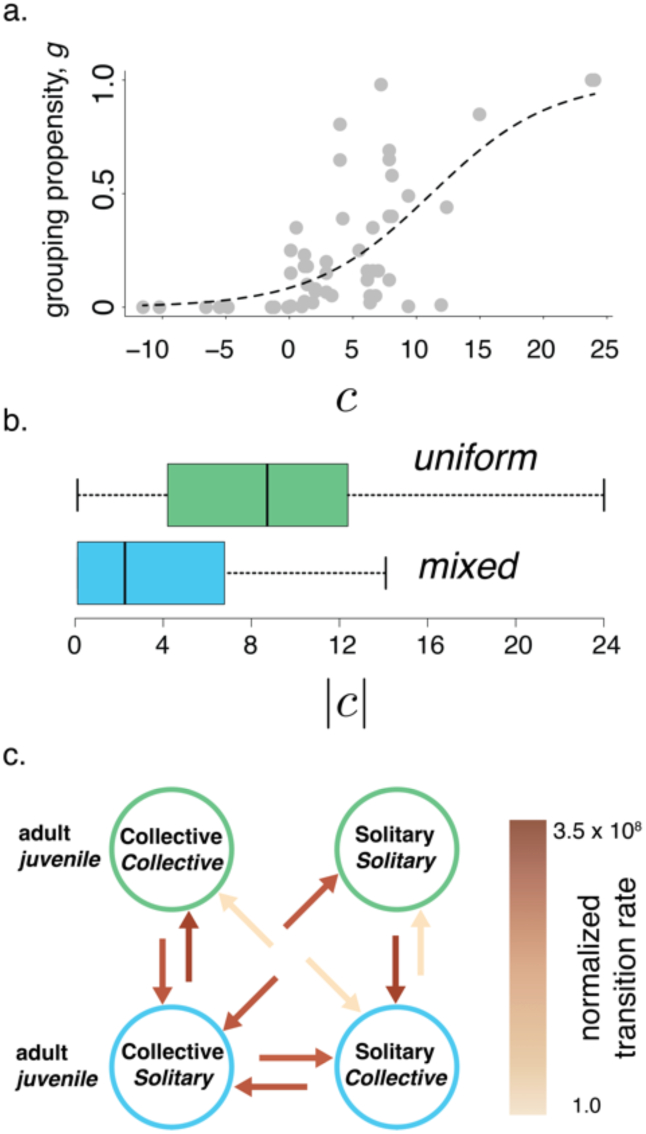
Grouping propensity and ontogenetic changes in the expression of collective phenotypes (a) Grouping propensity, *g*, the frequency with which species move collectively, as a function of latent collectivity, *c* (inferred from the analysis in Fig. 1b) for 71 species for which data on grouping propensity was available (see materials and methods). Curve is a logistic function fitted to data by least-squares. (b) Species that exhibit the same phenotype in pre-adult life stages as in adulthood (“uniform”, green) have higher values of |*c*| than do species that differ in their phenotype across life stages (“mixed”, blue) (c). Evolutionary transitions between collective and solitary phenotypes at the adult life stage pass through a phenotype with mixed life history (i.e., adult and juveniles have different phenotypes; blue circles); model selection supports a model that forces transitions between adult states to pass through mixed life history states by setting rate of direct transitions to zero.

### Variation in collective phenotype over ontogeny

Our analysis of grouping propensity suggests that species with mean values of *c* nearer the threshold are sometimes found moving collectively and sometimes found alone, but this analysis cannot discriminate whether this pattern is due to distinct subpopulations exhibiting different phenotypes, or whether individual animals may engage in collective movement facultatively. Some fish species are known to exhibit “mixed” life histories, in which individuals engage in collective movement at one life stage (e.g., larval, juvenile, adult) but not others. To more fully characterize mixed life histories, we gathered data on collective movement phenotype in juveniles and adults of the same species (See materials and methods). We found that the magnitude of collectivity, |*c*|, is significantly lower among species with mixed life history than among species with uniform life history (DF = 143, R^2^ = 0.21, *p* = 3.5 × 10^!$^, Fig. 3b), confirming that species nearer the threshold are more likely to exhibit both behaviors over their lifetimes. Interestingly, a similar fractions of species are solitary as juveniles and collective as adults (12% of the 145 species for which we had data on the phenotypes of both life stages) and collective as juveniles and solitary as adults (18% of 145 species).

**Figure 3.**
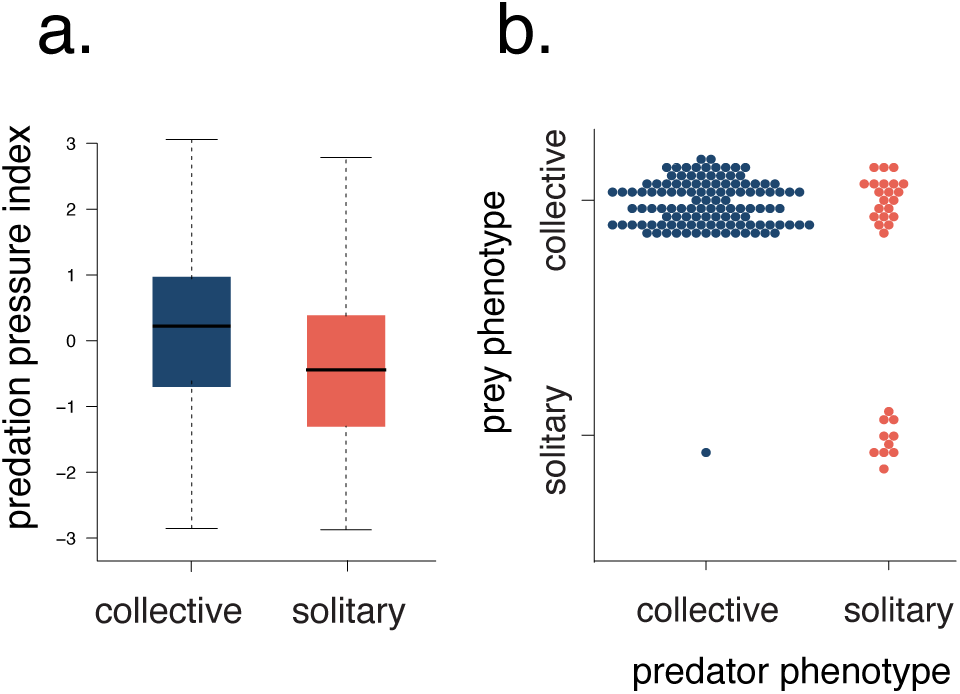
Ecological correlates of collective movement. (a) Collective species have, on average, higher values of predation pressure index than do solitary species. (b) Piscivorous species that exhibit collective phenotypes are more likely to eat other collective species than are solitary predators.

These findings imply that, on average, species with mixed life histories are nearer the threshold at which a species transitions between solitary and collective adult phenotypes. One explanation for this pattern is that modification of expression regulation in developmental pathways may be involved in the evolution and loss of collective movement in adults. This explanation is also consistent with recent within-species genomic and transcriptomic analyses that have implicated developmental genes in influencing engagement in collective movement (*49*). To more directly test whether evolutionary transitions in adult phenotypes involve intermediate stages in which adults and juveniles have different phenotypes, we performed an analysis of correlated evolution between mixed life history and adult collective phenotype. Using hidden Markov model selection (*50*), we found support for a model in which evolutionary transitions from one adult collective phenotype to another involve intermediate stages in which adult and juvenile phenotypes are mixed (Fig. 3; Δ AIC = 8.0 between models which assume no transitions between unform states [lowest AIC model, Fig 3c] and models in which all transitions are possible; See materials and methods).

Gene regulatory changes occurring early in the process of evolutionary divergence between closely related taxa have been implicated in facilitating phenotypic evolution, particularly for morphological traits (*51*, *52*). A related explanation for how traits can be labile in one lineage but stable in others involves ontogenetic shifts in gene regulation, which has been invoked to explain the evolution of inflorescence asymmetry in *Senecio* daisies (*53*) and tooth complexity in fishes (*54*). It may be that a similar explanation applies to our observations; species nearer the threshold *c =* 0 may express collective behavior facultatively in response to environmental conditions, which may promote expression at one life stage and not the other, while species with a higher magnitude of *c* may express their respective phenotypes obligately. Facultative expression is known to affect evolutionary elaboration and diversification of many traits (*55*). Following this elaboration, trait values can become less plastic due to consistent expression over generations of a reduced set of expression patterns triggered by the environment and the buildup of mutations in the regulatory pathway for unexpressed phenotypes, a process known as genetic assimilation (*56*).

### Ecological correlates of the evolution of collective movement

Collective movement appears to have evolved in many lineages, remained absent in others, and been repeatedly lost and regained in still others (Fig. 1b-d). Moreover, while some species are always found in collectives, others move collectively at some times and not others (Fig. 2a). Differences in the ecological factors that species are exposed to can drive facultative expression as well as divergent evolution. We therefore sought to determine whether ecological variables may be correlated with collective versus solitary phenotypes. In many animal lineages including fishes, collective movement is widely hypothesized to have evolved as an antipredator defense mechanism (*12*, *28*, *29*, *57*) (the “predation hypothesis”) although other selective forces such as locomotory efficiency (*24*), navigational benefits during migration (*29*, *58*), and the advantages of group foraging (*1*, *29*) have also been invoked. Our dataset allowed us to ask whether several of these hypotheses are consistent with data at a large phylogenetic scale.

The predation hypothesis holds that selection for predator avoidance can drive the evolution of collective movement (*12*, *29*). Through a number of mechanisms, collective movement can reduce predator-induced mortality experienced by animals that form collectives (*6*). To test the predation hypothesis, we sought to quantify predation pressure using an index that combined species trophic rank and body size. In general, large species that are high in the food web experience less predation pressure than small species that are positioned lower in the food web (*59*), and so we assume that species with higher index values experienced higher predation pressure than do species with low values (See materials and methods). If being exposed to high predation pressure leads to the evolution of collective movement, we expect that species with lower trophic rank and smaller body size (i.e., that have higher predation pressure index values) should be more likely to exhibit collective movement than species that occupy higher trophic ranks and have larger body sizes. Consistent with this, we found that collective species generally had higher values of estimated predation pressure (phylogenetically controlled analysis of variance, DF = 1706, F = 47.2, *p* = 0.0085; Fig. 3a, See materials and methods). Overall, the data suggest that species under more predation pressure are more likely to engage in collective movement (linear regression of predation pressure index as function of *c*: DF = 1706, R^2^ = 0.023, *p* = 2.0 × 10^−10^), consistent with the predation hypothesis.

A second hypothesis for the evolution of collective movement (the “collective foraging” hypothesis) posits that moving collectively enables animals to better exploit patchy ephemeral resources (*1*, *13*, *29*) due to the enhanced search efficiency and resource tracking that moving collectively enables (e.g., *1*, *60*, *61*). If patchy, dynamic resources select for collective movement, we expect collective species to be more likely than are solitary species to rely on such resources. Focusing our analysis on piscivorous species (See materials and methods), we estimated resource patchiness as the proportion of the diet composed of collective versus solitary fish species, because collective prey are an inherently patchier food resource compared to solitary prey. We found that collective predators overwhelmingly eat collective prey (97%, Fig. 4c). By contrast, solitary predator species consume collective prey at a frequency (76%) that is similar to the overall frequency of collective fish species on the phylogeny (79%). This pattern is robust to phylogenetic control (SI Section S4). These results are consistent with the hypothesis that the benefits of collective foraging when exploiting patchy resources may select for the evolution of collective movement.

**Figure 4.**
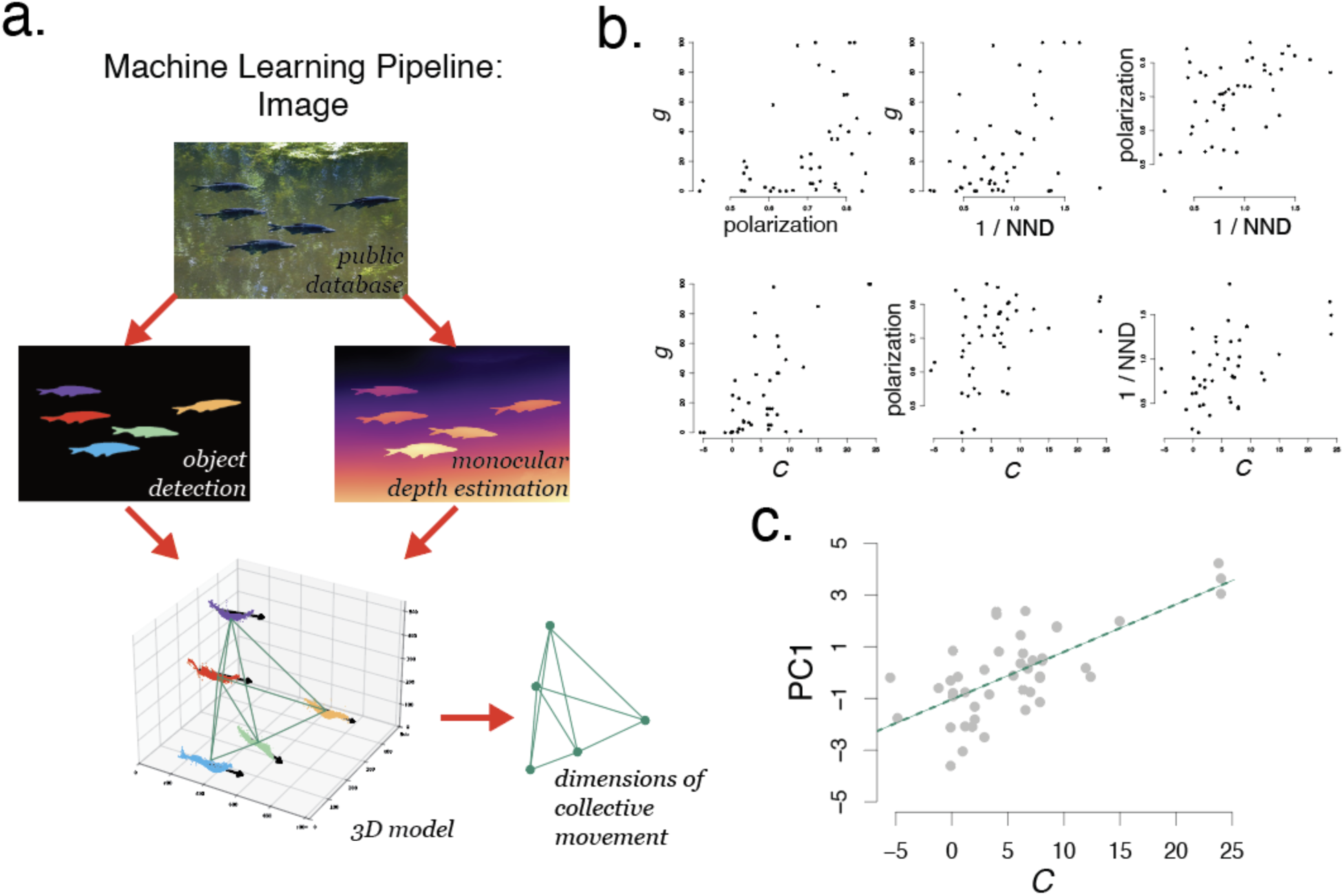
Continuous variation in collective movement phenotypes. (a) Machine learning pipeline for image processing (b) Component measures of quantitative features of collective movement are correlated with one another, and individual features covary with *c*, the hidden continuous trait inferred by the threshold (Fig 1a). NND: nearest neighbor distance (units of distance are body lengths); *g*: grouping propensity. (c) The primary axis of variation of measured components of collective movement (PC1, which explains 59% of the variation in the data), shows a strong relationship with *c* (DF = 43, R^2^ = 0.48, *p* = 9.9 × 10^!&^), such that higher levels of *c* are associated with a more strongly collective phenotype featuring larger, denser groups, higher alignment, and higher grouping propensity.

While high predation pressure and reliance on patchy resources are consistently correlated with the collective phenotype, there remains substantial variation in phenotype that is not explained by these ecological variables alone (e.g., Fig. 3a), suggesting that other factors also shape the pattern of diversity in collective movement. Indeed, species that engage in collective movement are found in many different habitats and across the globe: in freshwater (e.g., Cyprinid minnows) and saltwater (e.g., Engraulid anchovies), in benthic (schooling Sebastid rockfishes) and pelagic habitats (Scombrid tunas), and at high (e.g., Salmonids) and low (e.g., schooling Cichlids) latitudes. Combined with the observation that some large clades containing species that vary in ecology and life history are either exclusively solitary or exclusively collective (Fig. 1c), our results suggest that observed patterns of diversity in collective behavior are determined by a combination of facultative expression and vertical inheritance of trait values shaped by selection.

### Continuous variation in collective movement phenotypes

As discussed above, threshold traits are expressed as discrete phenotypes but are generally understood to be under highly polygenic control (*45*). We found that by treating collective movement as a threshold trait, we could produce trait value predictions that are consistent with the observed distribution of discrete phenotypes at a very large phylogenetic scale (Fig. 1a, compare tip states to branch colors; SI sections 1-2). However, past detailed analyses of collective movement behavior within species (*30*, *31*, *49*) and among small sets of species (*25*) have revealed what appears to be continuous variation in collective movement behavior. We therefore wondered whether the hidden variable *c*, inferred using binary phenotypes (which are frequently reported in the literature) might be correlated with measurable variation in the quantitative features of collective movement (which are far less commonly measured (*19*)) across species.

To answer this question, we sought to establish more detailed measures of collective movement in a subset of species spanning the phylogeny and varying in estimated value of *c.* To do this, we used large public image databases (iNaturalist [*62*], FathomNet [*63*]) containing imagery of fishes moving in groups in the wild. To make quantitative measurements from this imagery, we developed a computer vision pipeline that combined object detection and segmentation with state-of-the-art “monocular depth estimation” see materials and methods(*64*) (Fig. 4a). This allowed us to quantify relative positions and alignments of individuals within groups, variables that are widely used to quantify collective structure in animal groups (e.g., [*7*, *31*, *37*, *65*]). We combined these fine-scale measures of collective movement with the measure of grouping propensity described above, which yielded complete data for 45 species spanning 18 families.

Using this dataset, we compared the values of component measures of collective movement across species and found that these measures are often correlated with one another (Fig. 4b). A principal component analysis (PCA) of these trait data revealed that most trait variation is confined to a low-dimensional manifold (73% of variance is explained by PCs 1 and 2), and that trait variation within this low-dimensional space is highly correlated with *c* (Fig. 4c, linear regression between PC1 and *c*: DF = 43, R^2^ = 0.48, - = 9.9 × 10^!&^). To account for nonlinearity in the relationships among variables, we also performed an analysis of traits using a nonlinear dimension reduction technique called diffusion maps (*66*) as an alternative to PCA, which confirmed that variation in phenotypes among species is confined to a low-dimensional manifold (SI Section S5). As in the case of the PCA, phenotypic variation along the primary axis of this nonlinear manifold correlates strongly with a species’ collectivity, *c*, inferred from our phylogenetic analysis (linear regression: DF = 43, R^2^ = 0.45, - = 5.3 × 10^!’^).

These findings imply that species from lineages with a longer evolutionary history of moving collectively (i.e., those with higher *c*) also move collectively more frequently, form larger, denser groups, and are more aligned with neighbors. These findings appear to contrast with results from some within-species analyses of collective movement behavior, which have argued that different aspects of collective movement have modular genomic bases, which could enable them to evolve independently (*31*). At the scale of the cross-species comparison conducted here, traits associated with collective movement strongly covary.

### Collective movement is associated with increased macroevolutionary rates of diversification

The evolution of traits that strengthen barriers to reproduction can facilitate speciation(*67*, *68*). Collective movement not only involves the spatiotemporal clustering of individuals into distinct groups, but it can also induce the within-group synchronization of other life-history features such as migration and breeding phenology (*22*). Together, these features have the potential to increase spatial and allochronic reproductive isolation between grouping subpopulations (e.g., [*22*]) or lead to pre- or post-mating isolation as subtle phenotypic differences across groups are reinforced or grow through drift (*69*, *70*).

To determine if lineages in which collective movement has evolved also diversify more quickly, we compared alternative state-dependent diversification models, including models that allowed for hidden states (HiSSE [*71*]). This analysis revealed that net diversification rate was significantly higher in lineages expressing the collective phenotype than in those expressing the solitary phenotype (net diversification rate = 4.26 × 10^!(^ versus 3.03 × 10^!)^; trait-dependent model favored over best trait-independent model by 164 AIC points; SI Section S3, Table S1). To evaluate the robustness of this finding, we also used an alternative approach that jointly treats diversification and trait evolution as a reaction-advection-diffusion process (SI Section 3, Figure S3); this analysis also supports the conclusion that the extant phenotype distribution is consistent with a higher net diversification rate among collective lineages. These findings provide support for the hypothesis that engaging in collective behavior enhances diversification, either by increasing speciation rate, decreasing extinction rate, or both. Although the nature of our data limit our ability to infer the precise mechanism underlying this pattern, because of the combined influence of collective behavior on absolute fitness (*1*, *14*) and on spatiotemporal aggregation, we expect that the association between collective movement and increased diversification rate reported here may also occur in other clades such as mammals, birds, and insects.

## Discussion

In this work, we leveraged large public databases and novel machine learning methods to establish behavioral phenotypes for thousands of species of fishes spanning more than 270 families. This allowed us to study patterns of behavioral evolution across a vast phylogenetic scale. Our findings illustrate that collective movement has a rich and dynamic evolutionary history and that this collective phenotype is likely to have shaped the trajectory of phenotypic evolution and diversification within this largest of all vertebrate classes. Importantly, much past work on collective behavior has emphasized the fitness benefits it confers, but the frequent losses of collective movement documented in our study imply that there may often be selection against behaving collectively, for example due to increased intraspecific competition, targeting by specialized predators (Fig. 4b, [*72*]), increased risk of infectious disease (*73*), or informational costs (*74*, *75*).

More generally, our results provide three key observations that will aid in developing a fundamental understanding of the process by which collective behaviors and other collective phenotypes evolve. First, we show that collective movement exhibits a strong phylogenetic pattern (Fig. 1b), with clear evolutionary origins and subsequent diversification of clades that share the collective phenotype, as well as shared ecological correlates that are consistently associated with one phenotype or the other (Fig. 3). Similar patterns of clear heritability, large-scale phylogenetic structure, and common ecological correlates are found in numerous morphological traits that define phenotypes of individual organisms (e.g., [*51*, *53*, *54*, *76*, *77*]); our results show that these same patterns can be found in collective phenotypes. Second, by comparing classical measures of collective movement (*7*) across species (Fig. 4), we find continuous variation in the “strength” of collective movement that is confined to a surprisingly low-dimensional trait space. Across the range of species studied here, species with longer evolutionary histories of moving collectively also move collectively more frequently, group more densely, and align their motion more closely. Whether this pattern of strong correlations among the dimensions of collective movement is driven by correlated selection on these traits or by structural constraints imposed by the genomic architecture of these traits (or both) is presently unknown. Third, we find that clades that exhibit collective movement exhibit higher rates of macroevolutionary diversification than do solitary clades. To move collectively, individuals of a given species must be present in the same physical location at the same time. This requirement provides a clear mechanism that could lead to reproductive isolation of sub-populations, thereby promoting speciation. However, to our knowledge, the possible link between collective behavior and macroevolutionary diversification has never previously been tested.

Taken together, our findings reinforce results of microevolutionary work demonstrating that collective behavior is underpinned by heritable genetic variation (*31*, *34*, *65*), and that selective processes can drive evolution of these behaviors (*12*, *14*) and their elaboration once collective behavior originates within a lineage. We find support for classic hypotheses about the ecological factors that promote the evolution of collective movement, and we also show that collective behavior is frequently gained throughout the course of evolution but also frequently lost. Overall, our work establishes the importance of collective phenotypes in influencing the evolutionary history of vertebrates and points towards the potentially strong role such phenotypes may have played in shaping the broader diversity of life on Earth.

## Materials and methods

### Textual descriptions of behavioral phenotypes from FishBase database: Data processing and manual classification

We accessed the publicly available database, *FishBase v.*04/2025, which contains detailed data on fish species (47). Using the rfishbase API wrapper (79), we queried the “biology” table and extracted the “Comments” field for all species. The “Comments” field contains written descriptions of the biology of fish species contributed by experts and laypeople from around the world and includes in-text citations and full references. We searched these descriptions for a set of keywords: terms designed to include common terminology related to collective movement (school, aggregat-, shoal, solitary, alone, singl-, group, pair). We returned all records containing matches to one or more keywords and used this subset of all species records in FishBase as our preliminary text dataset.

Many records included in the preliminary dataset required manual removal because keywords were used in a context not relevant to the present study (e.g., “sandy shoals” or “targeted by solitary fishing vesselsℍ). We developed a simple user interface to display each record text with species identity hidden and keywords highlighted, allowing the efficient reading and manual classification of each record. Records for which class was uncertain were flagged for additional review. JL read each record, performing manual classification, and AG blindly validated flagged records and a subset of non-flagged records. The reviewer agreement rate was 100%. Phenotype classifications from this manual screening step were not used in any downstream analyses. Rather, we used this procedure to (1) identify a subset of FishBase entries that contained sufficient information about behavior to attempt to classify phenotype, and (2) verify that independent human experts could read this information and come to consistent conclusions regarding phenotype, further reinforcing the idea that text within the subset of FishBase entries we identified as candidates contained meaningful content related to collective phenotype.

### Building a ground truth dataset to train text classifier

Because there is the potential for error, ambiguity and inconsistency in the text descriptions within FishBase, we opted to identify phenotypes using a machine learning-driven text classifier, which provided both a consistent, repeatable mechanism of classification as well as a measure of uncertainty. To train such a classifier, we sought to generate a ground-truth data set containing high-confidence, expert-verified phenotype classifications for a subset of the FishBase text entries. Starting from a randomly generated, phylogenetically representative target sample of 600 taxa, we (JL, AG, EH, and AMH) conducted a search of literature and technical field identification guides that provided information on the behavior of each species. For each species, we assigned one of two classes: non-collective (“solitary”) or collective behavior. The collective class was therefore assigned to species who are described as exhibiting any behavior that involved aggregation into groups of >=3 individuals during regular activity, regardless of how coordinated the individuals in a group are described or assumed to be. The solitary class includes only species who have been verified to not form conspecific groups of more than two individuals. We did not include spawning or mating aggregations as evidence of engaging in collective movement, so mentions of grouping behavior under these conditions did not influence classification. We avoided using web resources that did not provide external citations for the behavior of interest, as those resources often used language similar to that found in FishBase (and we therefore assume that the resource was in fact drawing from FishBase to create their informational content). When the FishBase record provided a direct citation of a scientific article, report from a scientific organization, or a technical field identification guide for the behavior of interest, we did allow validation via that external citation when it was accessible and provided a clear description of the behavior of interest. JL reviewed all validation classifications and combined the resulting 329 externally validated records into a uniform ground truth dataset with citations (see SI Section S7).

### Assigning probabilities to discrete trait classifications using a machine learning classifier

To assign phenotypes and compute estimates of uncertainty for each assignment, we implemented a classification pipeline using ModernBERT-embed, an encoder-only transformer model optimized for text embeddings (*49, 80*). We fine-tuned this model using SetFit (Sentence Transformer Fine-tuning), a framework designed for few-shot classification that achieves high accuracy with limited labeled data (*48*).

SetFit operates in two stages. First, it fine-tunes the embedding model using contrastive learning on text pairs generated from the labeled examples, teaching the model to produce similar embeddings for texts within the same class and dissimilar embeddings for texts from different classes. Second, a logistic classification head is fitted on the resulting embeddings to output predicted labels (phenotypes in this case) for a given input text.

Using our ground truth dataset of 329 species records, we fine-tuned the model with a 50/25/25 train-validation-test split, stratified to preserve class distributions across splits. Prior to training and inference, we applied regex-based text cleaning to remove all in-text citations, HTML tags, and reference markers from behavioral descriptions. To address inconsistent performance with rare data types, we also augmented the training data split with paired, targeted synthetic texts prior to training (SI Section S6). The model classified species into two phenotype categories matching our ground truth labels: non-collective (“solitary”) and collective. We then applied the trained model to predict behavioral classifications for 2989 unlabeled species records, obtaining both predicted class labels and associated probability scores for each prediction.

The output of the model was a set of classification probabilities for each species in the prediction set. These probabilities were combined with the ground truth sets (for which binary probabilities of 0 or 1 were used) for use as data in the ancestral state reconstruction and tip-state inference, described below. All data and classifier models are available in the SI (SI Section S7).

### Phylogenetic comparative analyses

#### Generating the phylogenetic tree

We used a published phylogeny of Actinopterygian fishes (*50*) for our phylogenetic comparative analyses. The primary results were generated using a maximum clade credibility (MCC) tree produced from 100 sampled trees from the “all-taxon assembled” tree, pruned to include only species for which we had phenotype data. Because this method infers the phylogenetic relatedness of many taxa for whom molecular data was not available using only a taxonomy, using such a tree as the basis for analyses of trait evolution must take care that findings are robust to breaking assumptions made during the tree assembly^81,82^. For that reason, we repeated all of our methods on the RAxML ‘core’ tree, which is informed exclusively by molecular sequence data, therefore excluding many Actinopterygian species (SI Section S7). We also repeated all methods on 10 all-taxon sampled trees in order to account for potential artifacts induced by uncertainty in the MCC tree (SI Section S7). None of the qualitative conclusions presented in the main text were sensitive to these alternative assumptions.

#### Ancestral state reconstruction and continuous tip state estimation

To reconstruct the evolutionary history of collective movement and to better quantify the extant variation of the trait, we used a threshold model of trait evolution to infer ancestral node and extant tip states (*52, 54, 83*). This model assumes that discrete trait values are underlaid by a hidden continuously variable trait, often referred to as the ‘liability’, which we refer to as *collectivity,* (*c*). This variable is assumed to evolve via Brownian motion. To fit the model to our tree and trait probabilities for tip states, we used a discrete approximation of the threshold model, which employs a method to compute the likelihood (*84*) allowing the use of maximum likelihood to fit the model to data efficiently (*55*). We used this discrete approximation of the threshold model, as implemented in the *phytools* package in R (*85*). Input data consisted of a two-class probability distribution for each species and the maximum clade credibility tree, above. We used 400 bins to discretize the latent continuous trait space. We fit the threshold model to these data (using ‘fitThresh’ in *phytools*) and then simultaneously inferred the ancestral and extant states (using ‘ancr.fitThresh’ in *phytools*), recovering for each node and tip (1) probabilities of being in each discrete state and (2) probabilities of being in each discretized bin of the hidden continuous state, *l*, assumed by the threshold model. Taking the maximum value for each of these two probability distributions yields the maximum likelihood estimate (MLE) for (1) discrete and (2) hidden continuous trait values, respectively. To increase the efficiency of this computationally intensive task, we modified the ‘fitThresh’ function code by parallelizing the matrix exponentiation step prior to calculating the likelihood during model fitting.

#### Assessing collective behavior-dependent diversification

To determine if collective movement has influenced rates of lineage diversification in Actinopterygian fishes, we used a hidden state speciation and extinction model comparison approach (HiSSE; *hisse* package in R; [*74*]). We followed the standard approach (*51*) to fit trait-independent models with 0, 2, and 4 hidden states and a trait-dependent model with 4 hidden states. Model comparison was conducted by comparing model AIC values (*57*), and we evaluated the influence of collective movement on lineage diversification using the net diversification statistics (λ - μ) of the hidden states associated with solitary versus collective discrete character states in the model with the lowest AIC value. We evaluated the robustness of conclusions of this analysis using an independent approach based on formulating trait evolution and diversification as a reaction-advection-diffusion model (SI Section 3), which corroborated results of the HiSSE analysis.

### Quantifying continuous variation in collective movement phenotypes across species

#### Estimating grouping propensity

To estimate the frequency with which individuals of a given species move collectively, we compiled a set of data from 71 species for which quantitative estimates were available on the grouping propensity. This dataset was compiled from two sources. The first source was published field studies reporting data from surveys that quantified the frequency with which individuals of a given species are found in groups of three or more individuals versus alone or in groups of two. Because many species of fishes engage in courtship or aggressive interactions in pairs, we excluded cases where pairs of individuals were observed together because we could not determine whether these animals were moving through the environment collectively (e.g., engaging in foraging, navigation, predator evasion, etc.) or engaging in other kinds of pairwise interactions such as mating or dominance interactions. To identify candidate data sources, JL and AH searched the literature for field-based studies reporting either the proportion of time that individuals spent in groups or the incidence of detecting a group given the detection of an individual of a species during a transect or fixed-point occurrence survey. We extracted quantitative measures of grouping propensity from tables, plots, and electronic supplemental data of published, peer-reviewed studies that reported data matching our criteria (SI Section S7). When quantitative measures were not provided in text, tables, or supplemental data, but figures displayed the data graphically, we used a plot digitizer to extract quantitative data from the plot. The second source of data was unpublished field video surveys conducted by AGG, EH, MAG, and AMH. These surveys consisted of either moving video transect surveys or fixed point video sampling surveys conducted in Curaçao (Netherlands Antilles), Mo’orea (French Polynesia), or the Gulf of Thailand (Thailand). For each video survey, we quantified the proportion of observation time during which a species was solitary or in pairs versus in groups of three or more. We used the fraction of time spent in a group of three or more as a measure of grouping propensity.

#### Continuous measures of collective movement from imagery

Many studies of collective movement focus on proximate measures (*20, 44*) of the spatial organization and motion of groups. Doing this for more than a handful of species, particularly in the wild, has proven challenging in past studies due to the need for high quality imagery from which these variables can be quantified as well as the three-dimensional nature of many fish groups in the wild (*28*). To surmount this challenge, we developed a computer vision pipeline that sequentially combines three complementary deep learning approaches for extracting information from still images of fish groups. For each image, we manually drew a small number (1 to 2) of bounding boxes around example fish, which served as visual prompts for YOLO-E (You Only Look Once with Efficient prompting; [*86*]). The model then iteratively refined its detections across multiple passes, using previously detected bounding boxes as prompts for subsequent rounds, progressively identifying all individuals in the image with increasing confidence thresholds. Detected bounding boxes were passed to SAM2 (Segment Anything Model 2, [*87*]), which generated precise pixel-level segmentation masks for each fish, enabling accurate delineation of body boundaries even in densely packed groups. Finally, we applied Depth Anything V2 [*67*], a monocular depth estimation model that infers relative distances from a single 2D image using visual cues such as occlusion, relative size, and texture gradients. By overlaying segmentation masks onto the estimated depth map, we reconstructed the three-dimensional arrangement of individuals from still photographs (SI Section S7).

Depth estimates obtained from Depth Anything V2 provide relative depth up to an unknown global scale. To convert these estimates into a consistent three-dimensional representation, we rescaled the depth dimension by identifying a single multiplicative factor for each image that yields approximately isotropic spatial variation (i.e., comparable variance along the *x*, *y*, and *z* axes) in the reconstructed 3D point cloud. This yielded depth estimates in the same scale as the x and y coordinates. We used these rescaled depth maps as our z dimension in the downstream quantification of the dimensions of collective movement.

To gather imagery from a diversity of species, we used two large public image databases: *FathomNet* (*66*) and *iNaturalist* (*65*). Specifically, we conducted a search for images of aggregating or schooling behavior for the 71 species for which we had estimates of grouping propensity. Qualifying images clearly showed three or more individuals of the same species in a single photo (SI Section S7). When multiple images from the same iNaturalist observation met the criteria, we used the single clearest image of the set. We attempted to obtain 5 qualifying images per species, but fewer than 5 qualifying images were available for many species (e.g., those that are infrequently observed or do not often aggregate). Applying our computer vision pipeline, described above, to each qualifying image produced a set of *xyz* coordinates for each pixel belonging to each detected fish in units of fish body lengths. We then used these values to compute a number of commonly used measures of group size and collective structure. Specifically, the adjusted number of individuals in the group was quantified as the number of masks divided by the diagonal image dimension in body lengths. A centroid in the 3D surface was calculated for each mask, and then a matrix of pairwise Euclidean distances between mask centroids was generated. This distance matrix was used to calculate the nearest neighbor distances for each individual and the mean nearest neighbor distance of individuals within the group. Principal component analysis was used to acquire the major axis of variation for each mask (corresponding to the line segment aligned along the animal’s body axis in 3D), and the vectors of the first principal component were used to establish each animal’s heading direction, assuming the heading was aligned with the major body axis. We then generated a matrix containing the difference in heading angles of all pairs of individuals in the group, which we then used to compute nearest neighbor alignment values and the global polarization measure.

### Mixed and uniform life histories

Non-adult life stage behavior relevant to our study was reported within the species record for a portion of the species in our dataset. This allowed us to conduct a an analysis of the potential for changes in development to be correlated with the evolution of adult collective movement. Juvenile states were coded manually through a keyword search of the FishBase descriptions for taxa for which we had adult phenotype probabilities (see materials and methods, above) and subsequent manual review. Afterwards, species for which data on non-adult behavior were available were categorized into “uniform” or “mixed” behavior life history, where “uniform” taxa exhibit the same behavior in non-adult and adult stages and “mixed” taxa exhibit different behavior in the different life stages. This life history trait was then treated as a separate discrete trait, and tests of correlated trait evolution were conducted. First, Pagel’s test was conducted using ‘fitPagel’ in *phytools* on the data subset. Then, a model comparison approach was implemented in a hidden Markov modelling framework of binary character evolution (*corHMM* [*88*]). Four models were fit, differing in assumptions of (1) a single rate (no hidden states) versus (2) 2 rates (1 hidden state), and (3) no transitions occurring directly between different adult, uniform life history states versus (4) the relaxed assumption that transitions can occur directly between different adult, uniform states. This allows a test of the dependence of adult trait evolution on non-adult developmental changes. If models making assumption 3 fit the observed data better than models making assumption 4, then it suggests that a changing collective phenotype across ontogeny is associated with transitions between solitary and collective adult states. AIC scores were used for model comparison.

### Trophic level, body size, and diet data

We aimed to estimate predation pressure as broadly as possible, using a proxy index. To compose this index, we first extracted trophic level estimates for all taxa in our study and species-level diet composition data for a subset of piscivorous species. FishBase includes trophic level estimates for all species using either diet composition data or, when diet composition is not available, data from taxonomically related species for which diet data is available and then leveraging the strong relationship between body length and trophic level in fishes (*89*); full details of the algorithm to assign trophic level estimates for each species are provided at https://www.fishbase.se/manual/English/FishbaseThe_FOOD_ITEMS_table.htm). Because our goal was to make reliable proxy estimates of predation pressure as broadly as possible, we exclusively used trophic level estimates that were derived from diet data, subsetting our dataset for the analysis to only those taxa. For the species remaining in the dataset, we extracted body length measurements from FishBase. We then conducted PCA on the two variables (trophic level and body length), and the resulting first principle component (which explained 67% of the variance in the data) served as our proxy index of predation pressure. To determine if collective species face higher predation pressure than do solitary species, we performed phylogenetically controlled logistic regression.

To determine if collective species were more likely than solitary species to have a diet that is patchily distributed, we subset the full dataset to piscivorous fish for which species-level diet composition data were available in FishBase (*47*), TroPhish (*90*), or the California Current Trophic Database (*91*). Because collective species are, by definition, patchily distributed in the environment, we used prey phenotype (solitary versus collective) as a coarse measure of the degree to which predators rely on patchy prey resources. For the data subset, we used phylogenetically controlled logistic regression (*92*) to determine if collective predators were more likely to eat collective prey than were solitary predators and vice-versa.

Because predator diets are often composed of multiple prey species, we used a permutation test to arrive at a robust and phylogenetically-controlled statistical assessment of the relationship of interest. At each permutation (of 1000), a single prey species was randomly selected from the diet composition data for each predator species with sampling probability equal to the proportion of the diet composed by the species, a phylogenetically controlled logistic regression, and p-value was recorded. The distribution of p-values was then used to judge the robustness of the relationship (SI 4, Figure S4).

## Acknowledgements

Conversations with Liam Revell improved the quality of this manuscript.

## Author Contributions

JL, MG, AF, and AMH conceptualized the project. JL, AG, EH, MG, and AMH collected and processed the data. JL, AN, LWK, AF, MP and AMH developed the analytical methods and wrote the code. JL conducted the analyses. JL and AMH wrote the paper. All authors edited and improved the manuscript.

## Data, code, and materials availability

All data are available in the manuscript or the supplementary materials.

## Supplementary Information for

### Section S1: Latent collectivity, *c*, relates to phenotypic *isolation* and phenotypic *stability*

To quantify the observed clade-wise variation in evolutionary stability of collective movement, we calculated two values for each species. First, we calculated a model-free measure of phylogenetic *isolation* (See materials and methods). *Isolation* measures the degree to which a species is isolated within the phylogeny from species with differing phenotypes. We measure *isolation* as the difference between the branch length distance from a focal species to its nearest neighbor with a different phenotype and the branch length distance to its nearest neighbor with the same phenotype (Fig. S1a). *Isolation* is positively correlated with |*c*| (Fig. S1b; linear regression R^2^ = 0.68, DF = 2837, p << 0.0001).

In order to more explicitly quantify the evolutionary stability of a trait within a lineage, we then calculated for each species the quotient of the average phylogenetic distance to all inferred (by the threshold model) threshold crossings and the average phylogenetic distance to all nodes. Values of this measure, which we refer to as *stability*, greater than 1 indicate that a taxon is less closely related to threshold crossings than to non-crossings, on average, and a value of less than 1 indicates that a taxon is more closely related to crossings than to non-crossings. The stability also shows a strong relationship with |*c*| (Fig. S1c, R^2^ = 0.56, DF = 2837, *p* = 2.2 × 10^-16^). This pattern is not driven solely by taxa with very high stability; excluding species with stability >= 1.06 still reveals a strong relationship (R^2^ = 0.31, *p* = 2.2 × 10^-16^).

**Fig. S1.**
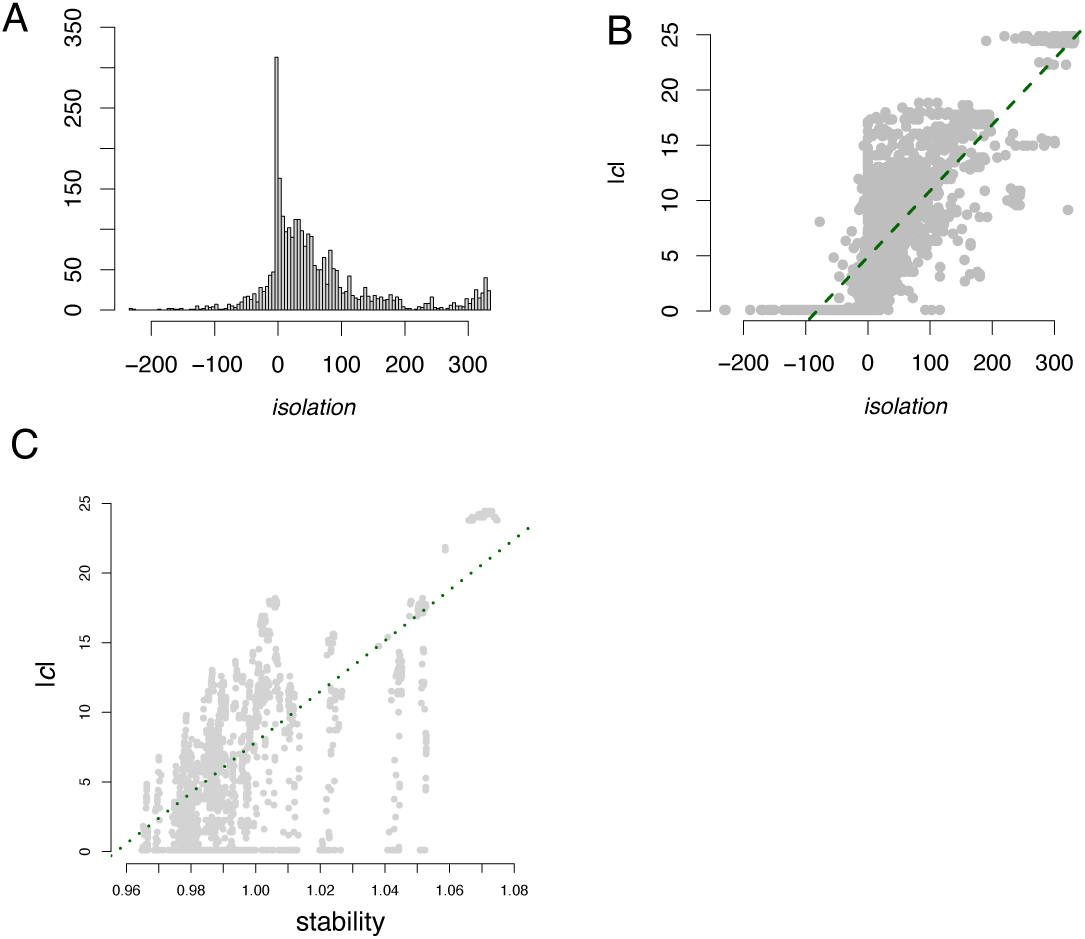
Trait *isolation* relates to how far removed a lineage is from a lineage with the opposite discrete phenotype. *Stability* relates to how far removed a lineage is from an ancestor with the opposite phenotype. (a) The distribution of *isolation* values for species. (b) *Isolation* is positively correlated with the magnitude of collectivity, |*c*|. (c) The magnitude of |*c*| inferred from the threshold model is strongly, positively correlated with *stability*.

### Section S2: Comparing evolutionary models of trait evolution for collective movement

The evolution of discrete traits is often modeled as a continuous-time Markov chain (*48*). These “Mk” models assume a constant probability of transitioning between discrete states throughout evolutionary time. While an Mk model can be fit to our data, we found that trait evolution simulations under the best-fit model (’fitMk’ in *phytools* [*85*]) on our phylogenetic tree did not recreate the distribution of *isolation* values present in the observed data (compare Fig. S2A and B; mean Kullback-Leibler divergence after 100 simulations for observed-simulated: 1.02; s.d. = 0.07). An alternative to the Mk model is the threshold model (see materials and methods). Under the threshold model, the rate of transitions from one discrete trait value to another need not be constant across a phylogenetic tree because transitions from one discrete value to another depend on the underlying value of continuous trait, *c* (*48, 44*). Given our finding that the rate of transitions from solitary to collective and *vice versa* appear to differ markedly across clades, we sought to test whether the threshold model could explain this pattern. Fitting a threshold model to our observed trait data (see materials and methods), we found that the distribution of *isolation* values closely matched that of the observed data (Fig S2c; KL divergence: 0.11). Indeed, comparing the fit of the Mk and threshold models to the observed data directly using AIC, the threshold model fits the observed data better than the Mk model (Mk log-likelihood: −1458.9, threshold log-likelihood: −1177.0, ΔAIC = 565).

**Fig S2:**
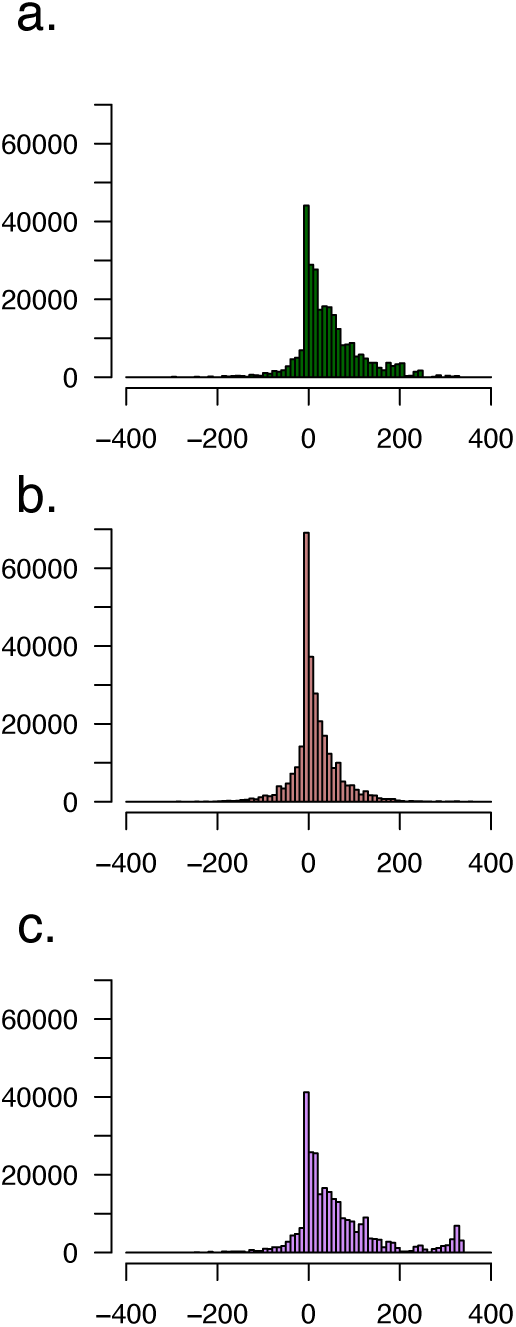
Distributions of the *isolation* values resulting from (A) the observed data, (B) 100 simulations on the best-fit Mk model, (C) 100 simulations on naive threshold models, (D) the best-fit threshold model. KL divergence values are shown in the supplemental text. Trait evolution under the Mk model (B) predicts that taxa will be less phenotypically isolated than that which is observed in the data (A).

### Section S3: An alternative model of trait evolution and trait-dependent diversification

In the Main Text, we report results from a hidden-Markov state-dependent diversification analysis (HiSSE), which shows that lineages in the collective state experienced an increased macroevolutionary diversification rate compared to lineages in the solitary state (Table S1). This approach is widely used in the field of macroevolution for this type of inference (e.g., [*54*]). To provide an independent test of whether observed patterns of trait evolution and lineage diversification are consistent with increased diversification in lineages that exhibit collective movement, we also used a second modeling approach. In particular, we developed a simplified model that jointly describes evolutionary diversification alongside evolution of collectivity, *c*. We note that the problem of evolution of a continuous latent phenotype *c* with the possibility of lineages leaving new species or going extinct can be described by a Fokker-Planck equation for a reaction-advection-diffusion model describing changes in the distribution of species with mean latent trait value, *c*:

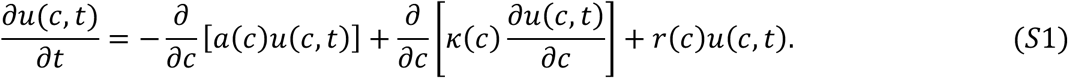

Here u(c,t) gives the frequency of species with mean trait value, *c*, at some time, *t*, since a common evolutionary reference point, k(c) describes how the distribution changes in response to unbiased increases and decreases in trait value (diffusion) a(c) describes directional changes in the distribution due to sustained, long-term directional selection, and r(c) describes the per-species rate of gain (or loss if r < 0) of new species. Note that we allow for the possibility that 4, 0, and 6 may vary depending on the value of). This latter assumption allows us to determine which combinations of *c*-dependent trait diffusion, directional selection, and diversification can produce patterns of latent trait values, *c*, that are consistent with those inferred from the threshold model fitted to our dataset.

**Table S1.** HiSSE model comparison. The trait-dependent, 4 hidden state model is favored over all character independent models. In the favored model, net diversification rate is much higher under collective (4.26 X 10^-2^) versus solitary (3.03 X 10^-3^) states. Net diversification calculated as λ - μ (see [*44*]).

| Model | AIC |
| --- | --- |
| Character Independent,<br>0 hidden states | 26594 |
| Character Independent,<br>1 hidden states | 26303 |
| Character Independent,<br>3 hidden states | 26071 |
| **Character Dependent,**<br>**3 hidden states** | **25906** |

To make inferences from Eq. (S1), we assumed an initial distribution of u(c,t= 0), given by a Gaussian distribution. For each term, k(c), a(c) and r(c) we considered both constant functions and sigmoid functions (with step functions as a limiting case), and allowed individual terms to be set to zero, enabling a wide range of possible dynamics involving unbiased changes in trait values (modeled by diffusion), persistent directional selection (modeled by advection) and lineage diversification (modeled by the “reaction” coefficient, r(c)). The partial differential equation (PDE) given by Equation (S1) was solved numerically using a standard PDE solver for sufficiently long times with boundary conditions, u(c = ∞, t) = u(c = −∞, t) = 0. For each parameter configuration, we evaluated whether the simulated dynamics produced a distribution u(c, t) that, at any time point, closely matched the inferred distribution of the latent variable *c* across phylogenetic tips. We denote the time associated with the best match, t*. Bayesian optimization (the surrogateopt algorithm implemented in MATLAB) was used to identify parameter configurations of the equation and initial condition that reproduce the inferred distribution of *c* (Fig. S3, middle panels).

**Fig. S3.**
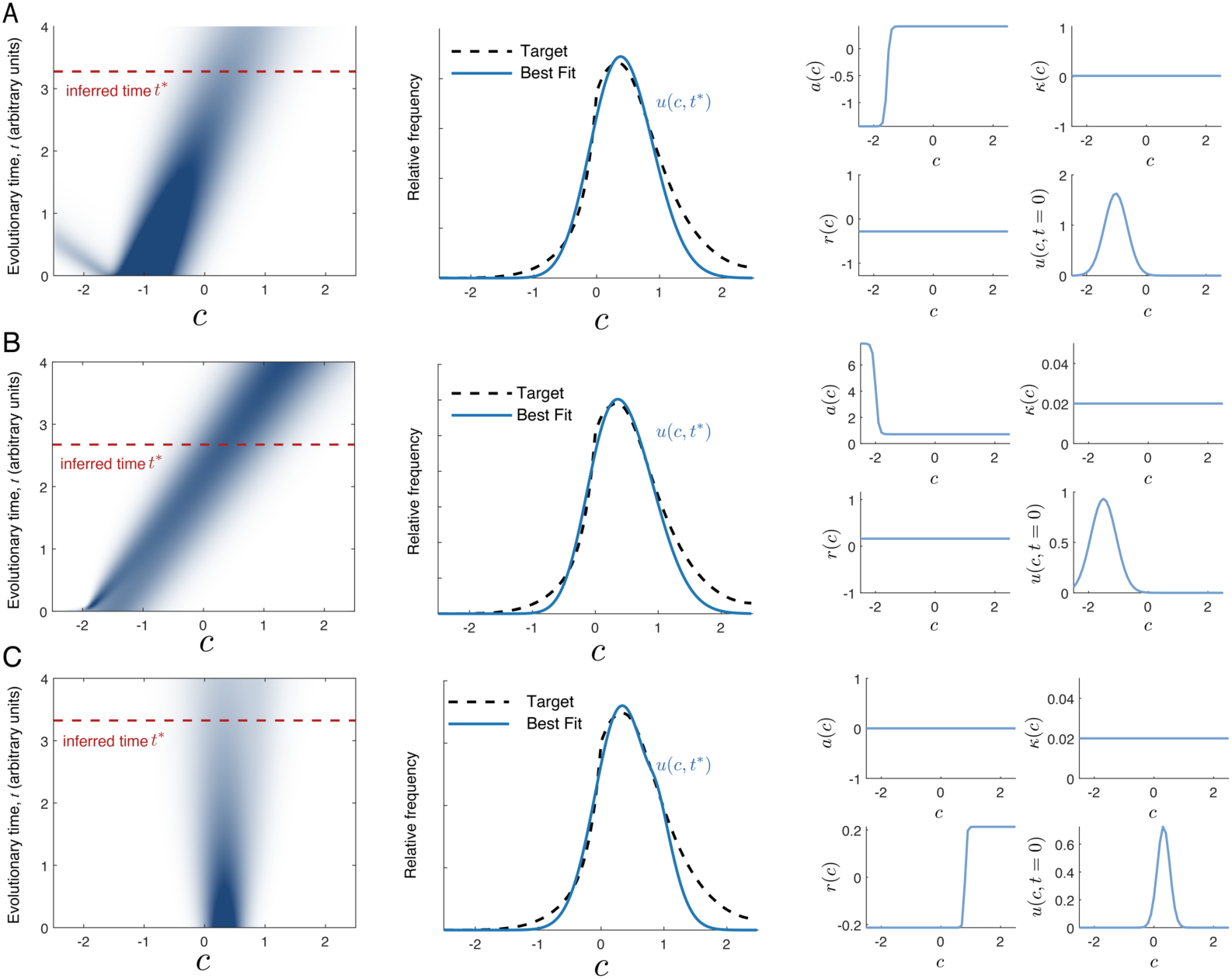
Results of Reaction-Advection-Diffusion model of trait evolution. (A-B) Example models in class one show highly non-stationary patterns in the distribution of *c* values across species over evolutionary time (left panels, darker shades correspond to a higher frequency of species with a given value of *c*, lighter shades correspond to lower frequency). Predicted distribution of *c* at time, t^∗^ (middle, blue curve) is consistent with empirical distribution inferred from threshold model analysis (middle, dashed curve). Trait dependent parameters in A and B involve strong, persistent directional selection indicated by non-zero values of a(c), along with uniform drift and uniform diversification. Under solutions in class two (example model shown in C), the distribution of trait values over evolutionary time is far more stable (left), leading to good agreement with the inferred distribution of *c* (middle), and evolutionary terms involve no sustained directional selection (i.e., a(c) = 0), but instead a constant diffusive term (k(c) = k > 0) representing unbiased increases and decreases in trait value which can be interpreted as resulting from temporally fluctuating selection or drift, and a trait dependent diversification term (r(c)) that is positive for c beyond some c^∗^ > 0 and negative for c below this value.

Given the flexibility of Equation S1, we do not expect to find a unique individual model that produces an unambiguously superior consistency with data. Rather, the fitting procedure identified a set of models that produced time-dependent distributions of tip *c* values closely resembling those inferred from the threshold model. This set contained two classes of models. In the first class of models, strong state-dependent directional selection on *c*, state-dependent diffusion, or strong state-dependent differences in diversification rate produced highly transient distributions that could nonetheless closely resemble inferred patterns of *c* during brief windows of evolutionary time (Fig. S3A-B). In these scenarios, the distribution of *c* in our dataset represents a highly transient pattern that could be observed in contemporary species but would be very different if traced into the past or forward into the future under the same conditions. In the second class of models, state-dependent diversification in which lineages with) > 0 exhibit higher diversification rates than lineages with) < 0 leads to stable distributions of *c* that resemble those inferred from the threshold analysis for our dataset (Fig. S3B). This second class of models is consistent with the finding of the HiSSE analysis that the distribution of trait values observed across species in our dataset is consistent with higher diversification rate within lineages expressing collective phenotypes relative to lineages expressing solitary phenotypes.

### Section S4: Phylogenetic control by permutation to accommodate variable diet

In our analysis of the degree to which predators that engage in collective movement exploit prey that engage in collective movement, we used phylogenetically controlled logistic regression to test for a relationship between predator and prey collective phenotypes and found that collective predators overwhelmingly eat collective prey, whereas the frequency with which solitary predators eat collective prey is roughly equal to the fraction of species in our dataset that exhibit the collective phenotype (Main; Fig. 3b). However, some species have diets that contain both collective and solitary prey species. We used the following approach to account for such mixed diets. For each predator species, we randomly selected a single prey species from the diet composition data for that predator with sampling probability equal to the proportion of the diet composed by the prey species. We then performed a phylogenetically controlled logistic regression in which prey phenotype (solitary or collective) was used to predict predator phenotype (solitary or collective). The p-value associated with the effect of prey phenotype on predator phenotype was recorded. This procedure was repeated 1,000 times using the sampling procedure described above to yield a distribution of p-values associated with the effect of prey phenotype on predator phenotype. The distribution of p-values exhibits a median of 0.007 and 90% of p-values fall below a significance level of 0.05 (Fig. S4).

**Figure S4.**
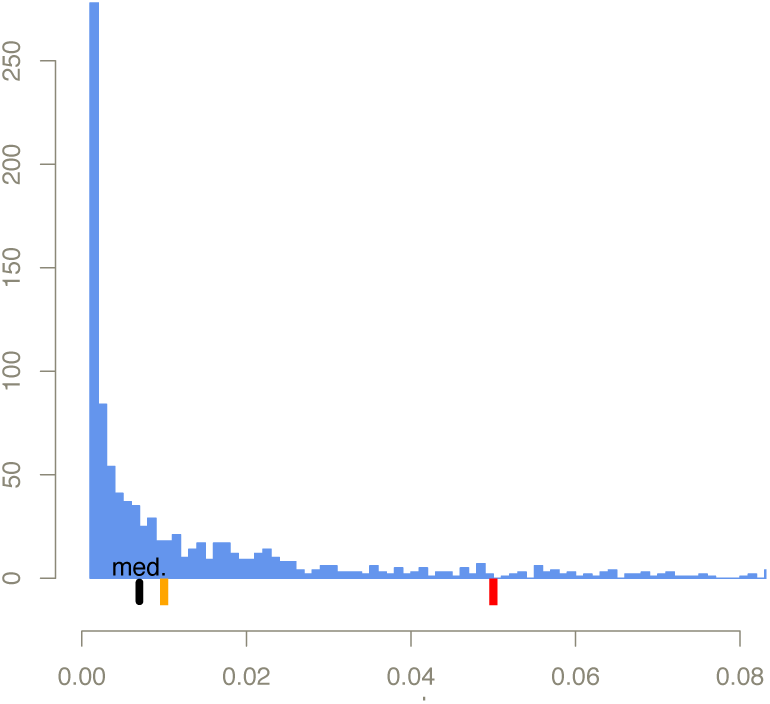
A phylogenetic control via permutation to test the relationship between predator and prey collective phenotype among piscivorous fish. Histogram bars show the frequency distribution of p-values from 1000 phylogenetically controlled logistic regression analyses on randomly sampled diet members. Rug ticks show confidence levels of 0.05 (red), 0.01 (orange), and the median of the distribution (black). N = 149 for all permutations. The results of this analysis suggest that collective predators differ from solitary predators in the phenotypic composition of their prey. See Main, Fig 3b.

### Section S5: Reducing the dimensions of collective movement

We employed two methods to identify primary axes of variation in quantitative traits related to collective movement. In the first method, we performed a principal components analysis (PCA) after centering all raw variables to a mean of zero and scaling them so that each variable had a standard deviation of one. This analysis indicated that 73% of the variance in these variables could be explained by variation along two orthogonal dimensions, principal component 1 (PC1, 59% of total variance) and 2 (PC2, 14% of total variance). Because there was some evidence of nonlinearity in pairwise relationships among raw variables we also performed a second analysis using a nonlinear manifold discovery method known as diffusion maps. Using this method, we characterized the geometry of quantitative trait space. Briefly, following column standardization we constructed a weighted k-nearest neighbors affinity matrix < from the species feature vectors, where the similarity between agents i and j was defined as the reciprocal of the Euclidean distance, 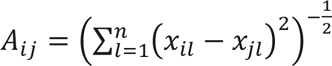 where x_il’_represents the *l-th* feature of observation *i*. We thresholded **A** to include only the K largest affinities for each data point and we then computed the row-normalized Laplacian, **L = I – D^-1^A**, where **I** is the identity matrix and **D** is the diagonal degree matrix with entries D = D_ij_ = ∑_j_ =A_ij_. The spectral properties of **L** reveal the intrinsic dimensionality of the data and the resulting eigenvectors, *ϕ*_k_, provide a coordinate system for the trait manifold whose orthogonal axes represent fundamental directions of variation in how species function in groups. Analysis of the Laplacian spectrum revealed a prominent spectral gap following the first non-trivial eigenvalue, suggesting that the variation in species’ quantitative collective traits is well-described by a one-dimensional manifold (*93–97*). Namely, we calculated the ratio of the first two non-trivial Laplacian eigenvalues, λ_3_/ λ_2 (_≈ 3.2, with λ_1_ = 0 as the trivial eigenvalue. This ratio indicates that the primary diffusion mode persists nearly three times longer than any subsequent mode suggesting a separation of scales. Further, Cheeger’s inequality 6 relates the first non-zero eigenvalue to the conductance, or bottleneck, of the affinity matrix **A** (*97–99*). The conductance h(G) is bounded by h(G) ≥ λ 2/2, and the observed gap between λ_2_and λ_3_ (λ_2_ ≈ 0.0011; λ_3_ ≈ 0.035) implies that the phenotypic variation is captured by a globally-consistent tradeoff. We therefore focused our analysis on the principal eigenvector, *ϕ*_1_, which reveals a major axis of variation where high values of *ϕ*I_1_ correspond to species that are often found in groups and form high density groups, that have large size, high local alignment, and low variances in these attributes, and low values along this axis imply the low values of these same variables and high variances.

### Section S6: Training data augmentation

Initial tests indicated that the model performed classification well overall, but was somewhat less reliable in certain settings (e.g., “conflicting information” deficiency, when records include language about population subgroups that behave differently, or “negative information” deficiency, when records include negative information such as “never forms conspecific groups” instead of positive information such as “is solitary”), resulting in reduced performance as quantified by the composite F1 score (SI 7). We therefore fortified the training data with a set of 32 synthetic records (SI 7 for synthetic texts used), increasing the training data split by 20%. In creating these text samples, we followed a paired approach by which two text samples were generated that contrasted in the class described, both using the same phrasing required to address the targeted deficiency (SI 7). Manual review of the outputs of fortified and non-fortified approaches showed that performance on rare data types (e.g., “negative information”) was improved by augmentation, and this was confirmed quantitatively by subjecting each model to a battery of synthetic texts designed to test the targeted deficiencies (SI 7). Importantly, the best-performing model was trained on a set that included augmentation and showed high performance (98% accuracy, 98% precision, 98% recall, and 98% F1 on held-out samples), an improvement over the non-augmented model version with the same data split (96% accuracy, 96% precision, 96% recall, and 96% F1), and we judged this model appropriate for use for our study.

### Section S7: Text classifier model and related supplementary files

The supplement contains a number of electronic materials related to our use of a text classifier model to process text descriptions of species’ behavior into binary trait data:

- “classifier_sensitivity.R” contains a script to interactively compile and evaluate reports of text classifier model fits (see SI Section S6). This is best run within an interactive R environment and allows for flexible exploration of the different classifier models that we evaluated.
- “./classifier_sensitivity/” is a directory containing .csv file outputs of text classifier model fits, which vary in the proportion of ground-truth data used in training and the degree of training data fortification used. These files are compiled into a more readable format using classifier_sensitivity.R, above.
- “./data_generation_pipeline/trainloop_f.py” is a python script that trains a series of text classifier models on different train/test splits and with different levels of synthetic training data fortification/augmentation. Users should be aware that the provided code requires specific computational resources (e.g., NVDIA graphical processing unit with sufficient memory).
- “fortification_text.csv” contains the synthetic texts used to augment/fortify the training data in order to address issues with classification of rare data types and sentiment sensitivity (see SI Section 7).
- “groundtruth_reference.csv” contains the records used for the groundtruth dataset, including references and details of review and quality control.

Other files not directly related to our use of a text classifier model:

- “README.txt” contains a guide to all materials in the electronic supplement, including those listed above and those used in the supplemental code.
- “pCollective_set.csv” contains the records for the estimation of grouping propensity, *g*, including references to sources used.
- “comp_data.csv” contains the primary dataset for analyses in “data_generation_pipeline/results_f.R”, downstream of the core data generation pipeline.
- “comp_data_core.csv” contains the primary dataset conducted on the ‘core’ tree (excluding taxa for which genetic information were unavailable during tree construction; see materials and methods) for analyses in “data_generation_pipeline/results_f.R”, downstream of the core data generation pipeline.
- “./dims/” contains the data for the dimensions of collective behavior analysis, with filenames indicating species name and public database accession number.
- “./etc/” is a directory containing other materials to reproduce analyses.

